# GREB1 regulates proliferation of estrogen receptor positive breast cancer through modulation of PI3K/Akt/mTOR signaling

**DOI:** 10.1101/704825

**Authors:** Corinne N. Haines, H.D. Klingensmith, Craig J. Burd

**Affiliations:** Department of Molecular Genetics, The Ohio State University, Columbus, Ohio, USA; The Ohio State University Comprehensive Cancer Center, Columbus, Ohio, USA

## Abstract

Over 70% of breast cancers express the estrogen receptor (ER) and depend on ER activity for survival and proliferation. While hormone therapies that target receptor activity are initially effective, patients invariably develop resistance which is often associated with activation of the PI3K/Akt/mTOR pathway. While the mechanism by which estrogen regulates proliferation is not fully understood, one gene target of ER, growth regulation by estrogen in breast cancer 1 (GREB1), is required for hormone-dependent proliferation. However, the molecular function by which GREB1 regulates proliferation is unknown. Herein, we validate that knockdown of GREB1 results in growth arrest and that exogenous GREB1 expression initiates oncogene-induced senescence, suggesting that an optimal level of GREB1 expression is necessary for proliferation of breast cancer cells. Under both of these conditions, GREB1 is able to regulate signaling through the PI3K/Akt/mTOR pathway. GREB1 acts intrinsically through PI3K to regulate PIP_3_ levels and Akt activity. Critically, growth suppression of estrogen-dependent breast cancer cells by GREB1 knockdown is rescued by expression of constitutively activated Akt. Together, these data identify a novel molecular function by which GREB1 regulates breast cancer proliferation through Akt activation and provides a mechanistic link between estrogen signaling and the PI3K pathway.

## Introduction

Breast cancer is the most frequently diagnosed malignancy in women (1). Over 70% of breast cancer patients are diagnosed with the estrogen receptor-positive (ER+) subtype, which is characterized by the expression of the transcription factor ER and dependence on ER activity for tumor cell growth and survival (2–6). Patients diagnosed with the ER+ subtype of breast cancer are typically prescribed endocrine therapies that target ER activity (2, 3, 6). However, resistance to endocrine therapies invariably occurs, leading to re-activation of the ER, expression of ER-target genes, and ultimately, patient relapse (2, 3, 6). Treatment options for patients that are resistant to endocrine therapies are limited, highlighting the need for innovative therapies that target downstream of the ER (2, 3, 6).

Crosstalk between the ER and the PI3K/Akt/mTOR pathway has been implicated in ER+ breast cancer progression and resistance to endocrine therapies (7–10). The *PIK3CA* gene, which encodes the catalytic subunit of the PI3K enzyme, is the most commonly mutated gene in ER+ breast cancer patients with over 40% incidence (11). This mutation is speculated to be a causal event in breast cancer progression, suggesting there is some need to upregulate this pathway in the development of ER+ breast cancer (11). Studies have indicated the ER can interact with PI3K to regulate its kinase activity in an estrogen-dependent manner (12). Further, inhibition of the PI3K/Akt/mTOR pathway increases ER expression and sensitivity of breast cancer cells to tamoxifen treatment, suggesting that activation of this pathway is associated with resistance to endocrine therapies through downregulation of ER (13). Together, these data indicate interdependence between ER and PI3K/Akt/mTOR signaling, however, the molecular basis and clinical relevance for this cooperation remains unclear.

Herein, we show that the ER gene target, growth regulation by estrogen in breast cancer 1 (GREB1), is a regulator of the PI3K/Akt/mTOR pathway, linking ER activation to this critical signaling pathway. Expression of GREB1 has been highly correlated to ER-positivity in breast cancer cell lines and patient samples (14–17). Previous studies have shown that knockdown of GREB1 results in significantly reduced proliferation and colony formation of ER+ breast cancer cell lines indicating, a required role for GREB1 in regulation of estrogen-dependent proliferation (16–18). However, it appears that an optimal level of GREB1 expression is necessary for proliferation of breast cancer cell lines as exogenous expression of GREB1 inhibits growth (18). Interestingly, growth repression by exogenous expression of GREB1 was also observed in ER-negative cell lines, indicating the ability of GREB1 to regulate proliferation of breast cancer cells is independent of ER activity (18). Despite the clear association of GREB1 and proliferation of ER+ breast cancer, the molecular function of the protein and the mechanism by which it regulates proliferation remain largely unknown. In this study, we characterize a novel mechanism by which GREB1 regulates proliferation of ER+ breast cancer cell lines through activation of Akt.

## Materials and methods

### Cell lines and reagents

MCF7, T47D, ZR-75-1, and HEK-293T cells were validated using Short Tandem Repeat analysis by the Genomics Core in the Research Technology Support Facility (Michigan State University, East Lansing, MI 48824). HCC1500 cells were purchased from the American Type Culture Collection. Cells were maintained as previously described (18). For experiments with EGF stimulation, cells were cultured in serum-free media for 16 hours before being stimulated with 1 ng/mL recombinant human EGF (Thermo) for the indicated time. The inhibitors GDC-0941 was obtained from Cayman Chemicals and were used at the indicated concentration for 24 hours prior to harvest of the cells.

### Plasmids

3XFLAG PCDNA and H2BGFP have been described previously (18–20). GIPZ lentiviral non-specific shRNA (# RHS4346) and lentiviral GREB1-targeted shRNA plasmids (V2LHS_139192 and V3LHS_372339) were obtained from Open Biosystems. MISSION shRNA constructs targeted to PIK3CA (TRCN0000196 582, TRCN0000195 203, TRCN0000010 406), PTEN (TRCN0000002 745, TRCN0000002 747, TRCN0000002 749), and PDK1 (TRCN0000001 476, TRCN0000039 778, TRCN0000039 782, TRCN0000010 413) were purchased from Sigma Aldrich. Myristoylated (Myr) AKT1 from pBabe-Puro-Myr-Flag-AKT1 (21) was cloned into a pLenti-hygro backbone to create pLenti hygro Myr FLAG AKT1 (CA AKT) using standard Gibson cloning (NEB).

### Immunoblot analysis and antibodies

Cell lysates were prepared, subjected to immunoblot analysis, and visualized on a LI-COR Odyssey system as previously described (18, 22). Immunoblots were probed with the following antibodies: GREB1 (abcam; ab72999 or CST; P76195 Clone 9C1), β-actin (Cell Signaling Technologies (CST); 3700), phosphor-p38 (CST; 9211S), p38 (CST; 9212), phosphor-MEK1/2 (CST; 9121), MEK1/2 (CST; 9122), phosphor-ERK1/2 (CST; 4370); ERK1/2 (CST; 4695), phospho-MKK3/6 (CST; 12280), phospho-MSK1 (CST; 9595), phospho-ATF2 (CST; 5112), phospho-HSP27 (CST; 9709), phospho-MAPKAPK2 (CST; 3007), phospho-PTEN (CST; 9551), PTEN (CST; 9556), phospho-PDK1 (CST; 3438), PDK1 (CST; 5662), phospho-Akt Thr308 (CST; 13038), phospho-Akt Ser473 (CST; 9271S), Akt (CST; 2920S), PP2A (CST; 2041T), phospho-GSK3β (CST; 5558), mTOR (CST; 2983), Rictor (CST; 2114), Raptor (CST; 2280), and p110α (CST; 4249T).

### Adenovirus

GREB1 adenovirus was purified as previously described (18). Ad5-CMV-eGFP adenovirus (Baylor College of Medicine Vector Development Labs, Houston, TX 77030) was used as a control.

### Alamar blue assay

Cells were treated with 0.04 g/L resazurin sodium salt in phosphate buffered saline (PBS) at 37°C for 1 hour. Fluorescence was measured on a BioTek Synergy microplate reader using a 540/35 excitation filter and a 590/20 emission filter. Data are depicted as mean fluorescence normalized to day 0 ± SD for each condition from 3 biological replicates. Statistical significance for alamar blue assays was determined using either a two-tailed Student’s t-test (exogenous GREB1 expression) or a one-way ANOVA with post-hoc Tukey’s HSD test (shRNA experiments).

### SA-β-gal staining

Cells transduced with GFP or GREB1 adenovirus were plated on poly-L-lysine coated coverslips. Cells were fixed and stained for SA-β-gal activity using the Senescence β-Galactosidase Staining Kit (CST #9860) as previously described (23).

### Conditioned media assay

MCF7 cells were transduced with either GFP or GREB1 adenovirus. After 24 hours, transduced cells were washed in PBS and fresh media added. The following day, media was collected from the transduced cells and centrifuged at 500 × g for 5 minutes to pellet any cellular debris. Target cells were washed twice with PBS prior to the addition of conditioned media. After 24 hours, all cells were harvested by scraping in cold PBS containing 10 nM calyculin A (Cell Signaling).

### Co-culture assay

MCF7 cells were transduced with adenovirus expressing either GFP or GREB1 (both adenovirus vectors express GFP). The following day, transduced cells were trypsinized and replated at a 1:1 ratio with un-transduced MCF7 cells. Cells were harvested after 24 hours by trypsinization and washed twice with cold PBS containing 10 nM calyculin A (Cell Signaling). Cells were sorted from both the adGFP and adGREB1 co-cultures using a Becton Dickinson FACSAria II cell sorter into GFP-positive (transduced) and GFP-negative (un-transduced) populations. Cell lysates were prepared from the sorted populations and immunoblot analysis was performed as described above.

### Immunofluorescence microscopy

Cells were plated on poly-L-lysine-coated coverslips. Following treatment, cells were fixed in 4% methanol-free formaldehyde (Thermo) diluted in PBS for 15 minutes at room temperature. Cells were then washed three times in PBS before permeabilization with 0.5% saponin, 1% BSA PBS solution at room temperature for 15 minutes. The cells were labeled with the indicated primary antibodies for 2 hours at room temperature in a humidified chamber. Coverslips were then washed three times in PBS before incubation with secondary antibodies (Alexa Fluor 555 goat anti-mouse and Alexa Fluor 555 goat anti-rabbit; Invitrogen) at room temperature for 1 hour in a humidified chamber, protected from light. Coverslips were mounted on microscope slides with VECTASHIELD Hard Set Mounting Medium with DAPI (Vector Laboratories). Images were obtained using a spinning disk confocal microscope (Ultra-VIEW VoX CSU-X1 system; Perkin Elmer) and analyzed using Velocity (Perkin Elmer).

### PIP_3_ quantification

MCF7 cells were transduced with either GFP or GREB1 adenovirus. 24 hours after transduction, cells were placed in serum-free, phenol-red-free DMEM for 16 hours. Lipids were extracted and PIP_3_ levels measured using the PIP_3_ Mass ELISA kit (Echelon K-2500s) according to the manufacturer’s instructions.

## Results

### GREB1 initiates cellular senescence

Our previous work has suggested that an optimal level of GREB1 expression is necessary for proliferation of breast cancer cell lines. This work showed that both GREB1 knockdown (18) and exogenous expression of GREB1 results in growth arrest (Fig. 1A-B)(18). We previously reported that exogenous expression of GREB1 did not induce apoptosis (18), thus we investigated the ability of GREB1 overexpression to initiate oncogene-induced cellular senescence. Two ER+ breast cancer cell lines, MCF7 and ZR-75-1 cells, were transduced with adenovirus expressing either GFP or GREB1. Following 7 days of exogenous GREB1 expression, the cells were fixed and stained for SA-β-galactosidase activity, a marker of cellular senescence (23, 24). Compared to GFP control cells, cells overexpressing GREB1 had a large, flattened morphology and characteristic blue staining associated with SA-β-galactosidase activity (Fig. 1C). These data suggest that exogenous GREB1 expression is able to induce cellular senescence to inhibit proliferation of breast cancer cell lines.

**Figure 1.**
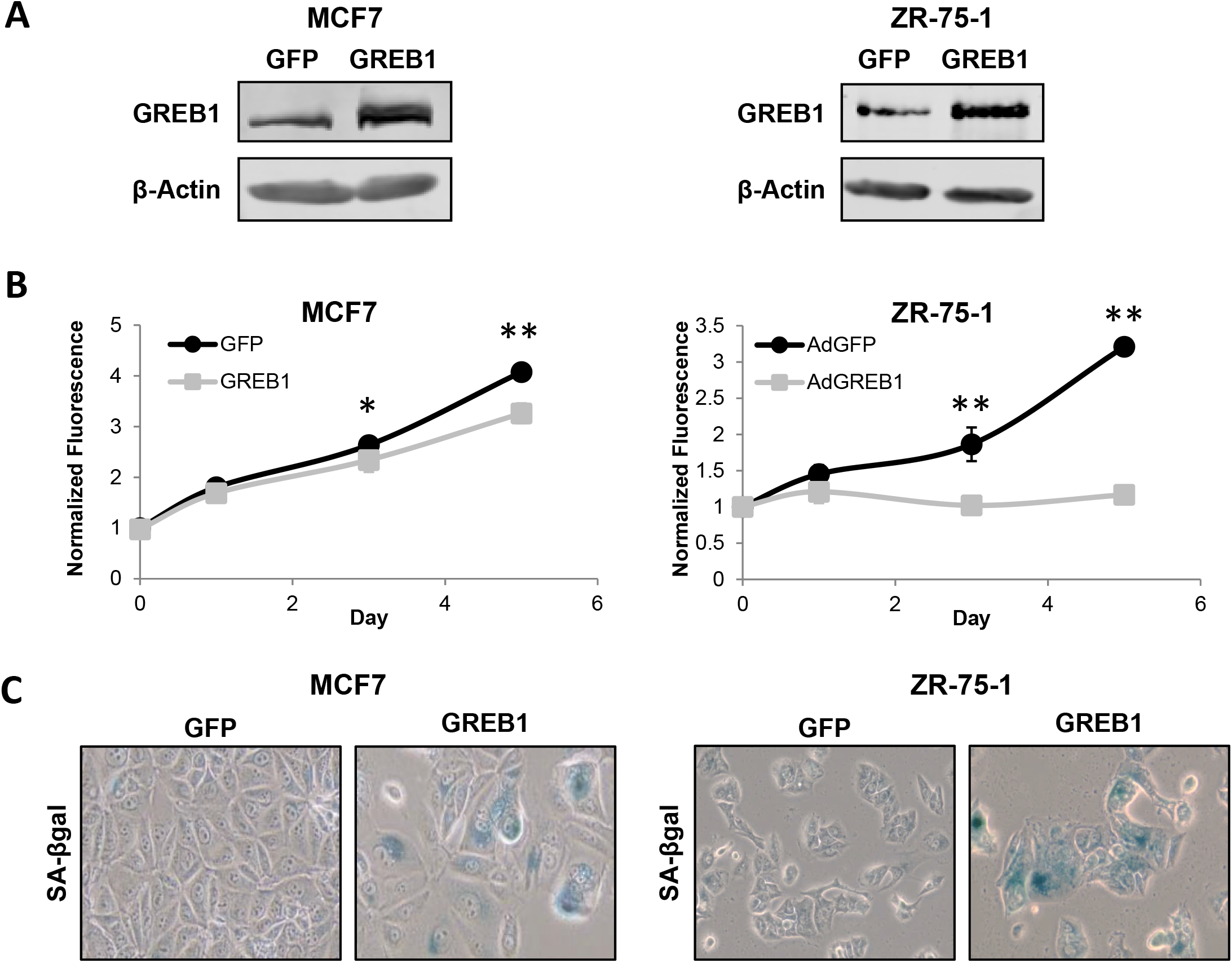
Exogenous GREB1 initiates cellular senescence. MCF7 or ZR-75-1 cells were transduced with adenovirus expressing GFP or GREB1. **A)** Immunoblot depicting the relative overexpression of GREB1 in cells transduced with GREB1 adenovirus compared to GFP control. **B)** Proliferation of transduced cells was measured by alamar blue assay. Data are plotted as mean fluorescence normalized to Day 0 ± SD; n=3 for each cell line. *p≤0.05, ** p≤0.005. **C)** Cells were fixed and stained for SA-β-galactosidase activity 7 days post-transduction.

### Exogenous GREB1 expression induces hyperactivation of the PI3K/Akt/mTOR pathway

In order to delineate the mechanism by which GREB1 regulates proliferation, we chose to focus our attention on two signaling pathways thought to play a critical role in regulating both senescence and proliferation phenotypes: the p38 MAPK pathway and the PI3K/Akt/mTOR pathway (25–27). MCF7 cells were transduced with adenovirus expressing GFP or GREB1 and after 48 hours, cell lysates were analyzed by immunoblot for activation of various nodes in the p38 MAPK pathway (Fig. 2A) and the PI3K/Akt/mTOR pathway (Fig. 2B). Our data indicate that exogenous GREB1 expression induces an increase in the activation and phosphorylation of p38 and its downstream effector, MAPKAPK2, however, the activation and/or expression of upstream regulators of p38 (MEK1/2, ERK1/2, MKK3/6) and other downstream effectors (MSK1, ATF2, and HSP27) were largely unaffected (Fig. 2A). Analysis of the PI3K/Akt/mTOR pathway revealed hyperactivation of Akt, as well as increased phosphorylation of GSK3β, a downstream effector of Akt, when GREB1 was exogenously expressed in MCF7 cells (Fig. 2B). As the PI3K/Akt/mTOR pathway is frequently altered in breast cancer, we analyzed the effect of GREB1 overexpression on Akt activation in a panel of ER+ breast cancer cell lines with wildtype or differing alterations to the PI3K/Akt/mTOR pathway (28). While MCF7, ZR-75-1, and T47D cells harbor mutations in this pathway, each cell line has varying levels of Akt activation with MCF7 cells demonstrating the least constitutive activity (Supplemental Fig. 1). In cells lines without constitutive maximal activity of this pathway, GREB1 induces a strong activation of Akt (Fig. 2C). In contrast, T47D cells, which have a PIK3CA mutation resulting in constitutively active Akt that does not respond to serum deprivation (Supplemental Fig. 1), are unaffected by GREB1 expression (Fig. 2C).

**Figure 2.**
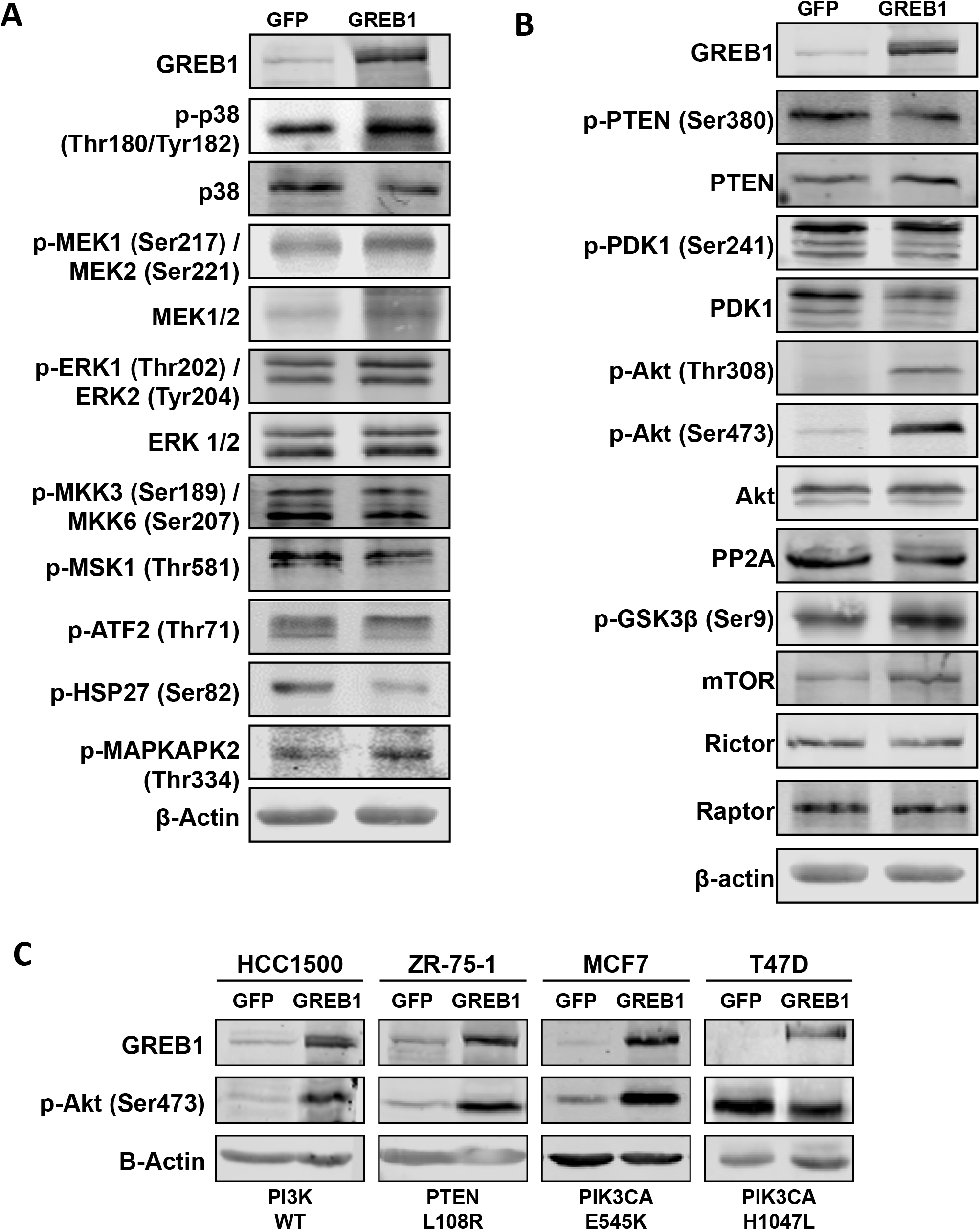
GREB1 modulates PI3K/Akt pathway signaling. A) MCF7 cells were transduced with adenovirus expressing GFP or GREB1. Cell lysates were harvested 2 days post-transduction and analyzed by immunoblot for indicated proteins in the p38 MAPK pathway. **B)** MCF7 cells were transduced with adenovirus expressing GFP or GREB1. Cell lysates were harvested 2 days post-transduction and analyzed by immunoblot for indicated proteins in the PI3K/Akt pathway. **C)** HCC1500, ZR-75-1, MCF7, and T47D cells were transduced with adenovirus expressing GFP or GREB1. Cell lysates were harvested 2 days post-transduction and analyzed by immunoblot for indicated proteins.

### GREB1-induced hyperactivation of Akt is PI3K-dependent

Akt requires phosphorylation at two sites, Thr308 and Ser473, by PDK1 and mTORC2 respectively, for maximal activation (29–31). Both phosphorylation events are dependent on PI3K and occur downstream of PI3K conversion of PIP_2_ to PIP_3_ (29, 31). Thus, we sought to determine if GREB1 was acting to regulate Akt activation upstream or downstream of PI3K. To this end, we targeted PI3K activity with the pharmaceutical inhibitor GDC0941. MCF7 cells were simultaneously treated with DMSO or GDC0941 and transduced with adenovirus expressing GFP or GREB1. After 24 hours, cell lysates were harvested and activation of Akt at Thr308 and Ser473 were evaluated by immunoblot analysis. The expected hyperactivation of Akt was observed in DMSO-treated cells that were transduced with GREB1 adenovirus (Fig. 3A). PI3K inhibition by GDC0941 demonstrated a marked decrease in basal Akt activation in the control GFP-transduced cells (Fig. 3A). A decrease in GREB1-mediated activation was also observed, particularly at Ser473 (Fig. 3A), suggesting that GREB1 is activating the canonical PI3K/Akt axis.

**Figure 3.**
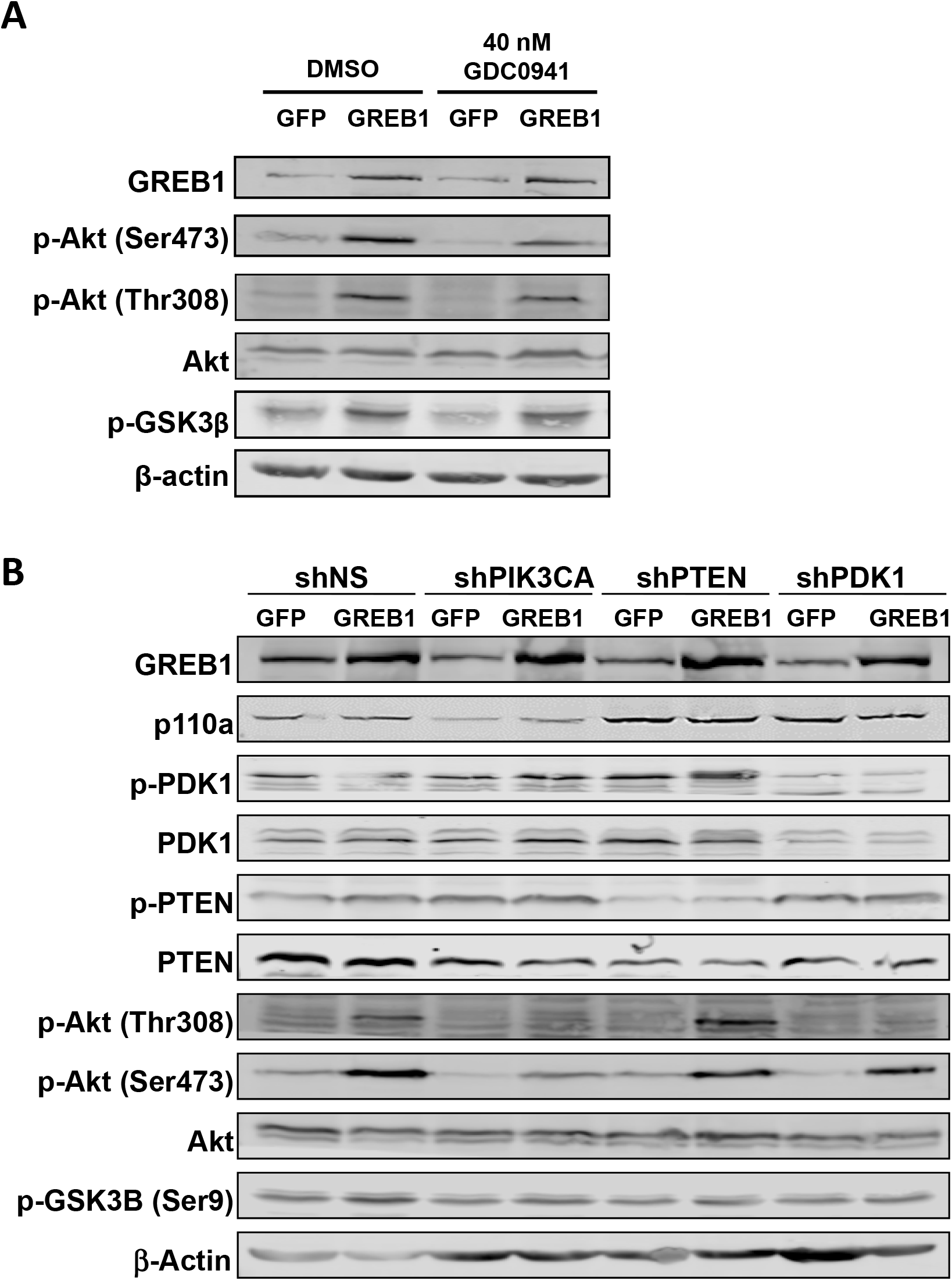
GREB1-induced hyperactivation of Akt is PI3K-dependent. **A)** MCF7 cells were transduced with adenovirus expressing GFP or GREB1 and treated with DMSO or 40 nM GDC0941 simultaneously. Cell lysates were harvested 24 hours post-transduction and immunoblot analysis was performed with the indicated antibodies. **B)** MCF7 cells were transduced with lentivirus targeted to a nonspecific control (shNS), PIK3CA, PTEN, or PDK1. Following selection, cells were transduced with GFP or GREB1 adenovirus. Cell lysates were harvested 24 hours post adenovirus transduction. Following SDS-PAGE, immunoblot analysis was performed with indicated antibodies.

To confirm this result and further probe other nodes in the pathway, MCF7 cells were transduced with lentivirus expressing shRNA targeted to a non-specific control (shNS), PIK3CA, PTEN, or PDK1. Following selection, cells were transduced with adenovirus expressing GFP or GREB1. After 24 hours, cell lysates were harvested and analyzed via immunoblot. Control cells transduced with non-specific shRNA demonstrated the expected increase in phosphorylation of Akt (Thr308 and Ser473) and GSK3β when GREB1 was exogenously expressed (Fig. 3B). Knockdown of PIK3CA expression reduced Akt activation in both control and GREB1-expressing cells (Fig. 3B). GREB1-induced hyper-phosphorylation of GSK3β was also reduced when PIK3CA was knocked-down (Fig. 3B). Knockdown of PTEN enhanced GREB1-induced hyperactivation of Akt at Thr308, suggesting the mechanism of GREB1 action is not through phosphatase inhibition (Fig. 3B). In contrast to our pharmacological approach (Fig. 2B), GREB1-induced hyperactivation of Akt at Thr308 was completely blocked by knockdown of PDK1, the primary kinase for this site (29–32)(Fig. 3B). As expected, PDK1 knockdown had no effect on GREB1-induced hyperactivation of Akt at Ser473, as this is not the primary kinase for this site (29–31) (Fig. 3B). Knockdown of PDK1 diminished GREB1-induced phosphorylation of GSK3β (Fig. 3B). These data further demonstrate that GREB1-induced hyperactivation of Akt is dependent on signaling through the canonical PI3K pathway.

### GREB1 activates Akt through intracellular mechanisms

Canonical activation of PI3K and Akt occurs through activation of receptor tyrosine kinases (RTK) or G-protein-coupled receptors (GPCR) by external stimuli (29, 33). We first sought to determine if GREB1-mediated Akt regulation is dependent upon induction and secretion of a signaling molecule that activates the PI3K/Akt/mTOR pathway. To this end, MCF7 cells were transduced with adenovirus expressing GFP or GREB1. Media from transduced cells was transferred to un-transduced MCF7 cells. After 24 hours, cell lysates were harvested and activation of Akt was analyzed by immunoblot. As expected, exogenous expression of GREB1 induced hyperactivation of Akt at Ser473 when compared to GFP-transduced cells (Fig, 4A). However, conditioned media was unable to induce hyperactivation of Akt in un-transduced cells (Fig. 4A).

**Figure 4.**
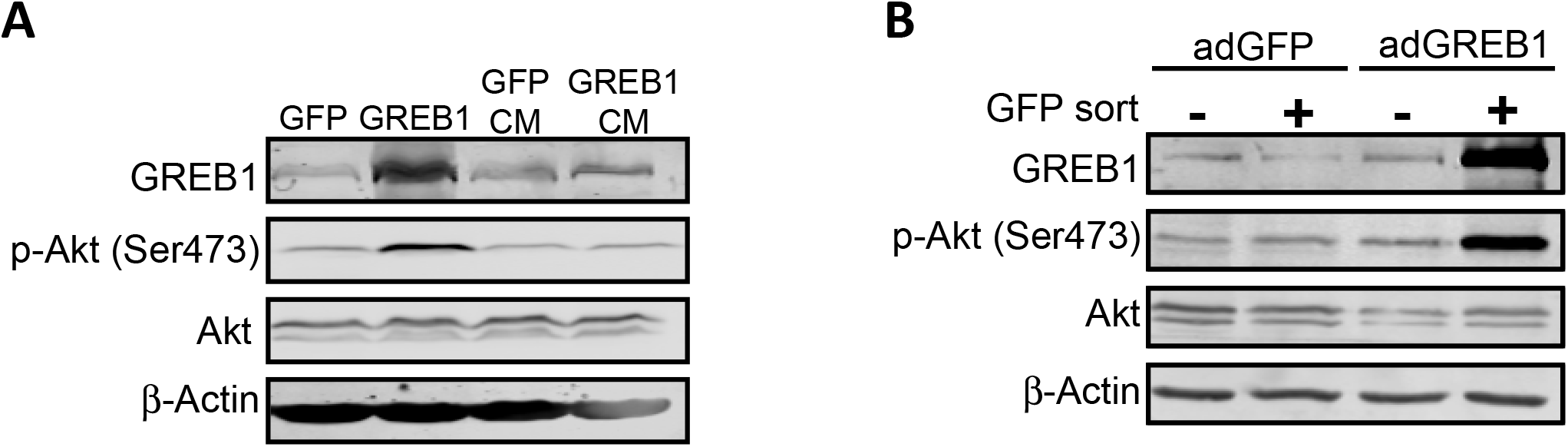
GREB1 activates Akt through intracellular mechanisms. **A)** Conditioned media from MCF7 cells transduced with GFP or GREB1 adenovirus was added to un-transduced cells. Cell lysates from transduced cells (GFP or GREB1) and un-transduced cells cultured in conditioned media (GFP CM or GREB1 CM) were harvested 24 hours later. Lysates were analyzed by immunoblot for indicated proteins. **B)** MCF7 cells were transduced with adenovirus expressing GFP or GREB1. Transduced cells were then cultured with un-transduced cells at a 1:1 ratio for 24 hours. Cells were harvested and sorted for GFP. Cell lysates were analyzed via immunoblot for expression of the indicated proteins.

Alternatively, exogenous GREB1 may induce the expression of a membrane-bound signaling molecule that could activate the PI3K/Akt/mTOR pathway. To investigate this possibility, MCF7 cells were transduced with adenovirus expressing GFP or GREB1 and then co-cultured at a 1:1 ratio with un-transduced MCF7 cells. After 24 hours, the cells were harvested and GFP-positive, adenovirus-transduced cells (GFP or GREB1), were sorted from GFP-negative, un-transduced cells. All populations were then analyzed for activation of Akt by immunoblot. Both GFP-transduced cells and cells co-cultured with the GFP-transduced cells had similar levels of Akt activation (Fig. 4B). Exogenous expression of GREB1 induced the expected hyperactivation of Akt at Ser473 within the transduced cells; however, the untransduced co-culture cells did not demonstrate Akt hyperactivation (Fig. 4B). While these data are negative, they clearly demonstrate that GREB1 regulates Akt activation through in an intracellular mechanism that does not require extracellular activation of RTKs.

### Exogenous GREB1 promotes recruitment of Akt to the plasma membrane

Akt is typically activated via recruitment to the plasma membrane by interaction with PIP_3_, however, it is believed that there are other pools of activated Akt on endomembrane surfaces and within the nucleus (29). To determine the localization of activated Akt induced by exogenous GREB1 expression, MCF7 cells were transduced with either GFP or GREB1 adenovirus. Following transduction, the cells were serum-starved for 16 hours to reduce basal Akt activation before stimulation of the pathway with EGF. Cells were then fixed and stained with DAPI and the indicated Akt antibodies. As both adenoviral vectors expressed GFP, we focused our imaging on transduced cell populations. In serum-starved, GFP-transduced cells, staining for activated Akt was minimal and staining for total Akt resulted in diffuse staining throughout the cytoplasm (Fig. 5A, Supplemental Fig. 2A-B). In contrast, cells transduced with GREB1 adenovirus under serum-starved conditions had distinct staining for activated and total Akt, primarily localized to the plasma membrane (Fig. 5A, Supplemental Fig. 2A-B). When stimulated with EGF for 5 minutes, both GFP- and GREB1-transduced cells demonstrated activated Akt and total Akt at the plasma membrane (Fig. 5A, Supplemental Fig. 2A-B). Activation of Akt and focal localization of total Akt at the plasma membrane was noticeably stronger in GREB1-transduced cells when compared to GFP-transduced cells in the presence of EGF (Fig. 5A, Supplemental Fig. 2A-B). These data further suggest that GREB1 may act through PI3K to increase PIP_3_ levels and Akt re-localization to the plasma membrane.

**Figure 5.**
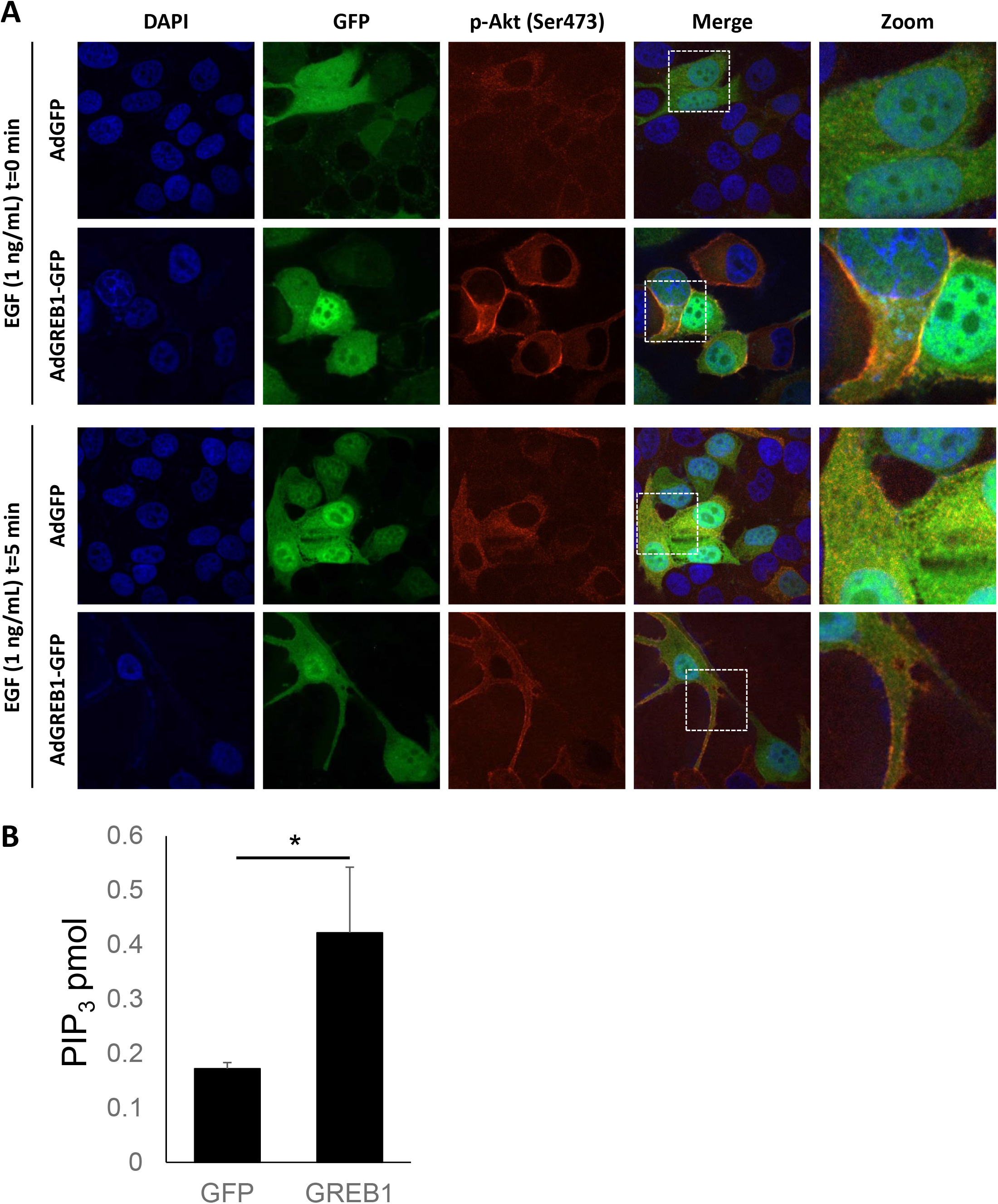
Exogenous GREB1 promotes recruitment of Akt to the plasma membrane. **A)** MCF7 cells were transduced with adenovirus expressing GFP or GREB1. The cells were then cultured in serum-free media for 16 hours before being stimulated with 1 ng/mL EGF for 0 or 5 minutes. Cells were fixed and stained for DAPI or p-Akt (Ser473). Immunofluorescence microscopy was used to visualize the activation and localization of Akt. **B)** MCF7 cells were transduced with adenovirus to express exogenous GFP or GREB1 and serum starved for 16 hours. Lipids were extracted from all samples and levels of PIP_3_ were measured via ELISA. Graphs represent mean PIP_3_ (pmol) + SD (n=3).

We sought to directly test if GREB1 expression influenced the conversion of PIP_2_ to PIP_3_. MCF7 cells were transduced with GFP and GREB1 adenovirus and then placed in serum-free media as depicted in Figure 5A. Following serum starvation for 16 hours, lipids were extracted and PIP_3_ levels measured by ELISA. Expression of exogenous GREB1 induced a significant increase in PIP_3_ levels, further indicating that GREB1 augments PI3K activity.

Previous studies have suggested that GREB1 is primarily localized to the nucleus in patient samples and breast cancer cell lines (15, 16), thus, it remained unclear how GREB1 was able to regulate signaling through a primarily cytoplasmic pathway. Interestingly, we discovered that under serum-starved conditions, endogenous GREB1 is diffuse throughout the cytoplasm and nucleus in MCF7 cells, but upon stimulation with EGF, the vast majority of GREB1 rapidly re-localizes to the cytoplasm (Supplemental Fig. 3A). As this is contradictory to previously published reports, we performed nuclear/cytoplasmic fractionation to verify cytoplasmic expression of GREB1. Under normal growth conditions (i.e. media containing FBS), GREB1 is primarily located within the cytoplasm of MCF7 cells (Supplemental Fig. 3B-C). Thus, in response to growth factor activation, GREB1 localizes to the cytoplasm wherein it can modulate the PI3K pathway.

### GREB1 regulates breast cancer proliferation through activation of the PI3K/Akt/mTOR pathway

In order to determine if GREB1-mediated Akt activation is imperative for proliferation of ER+ breast cancer cells, we tested whether constitutively activated Akt can rescue proliferation loss in GREB1 depleted cells. Thus, we made use of T47D cells which are ER+ and GREB1-expressing, but harbor a *PI3KCA*^H1047L^ mutation rendering this pathway constitutively active and unresponsive to typical PI3K/Akt/mTOR-activating stimuli, including GREB1 exogenous expression (Fig. 2C and Supplemental Fig. 1). T47D cells were transduced with shRNA targeting a nonspecific control (shNS) or shRNA targeting GREB1 (shGREB1 #1 or shGREB1 #2). Immunoblot analysis confirmed knockdown of GREB1, as well as hyperactivation of Akt in T47D cells (Fig. 6A). Proliferation of these cells was then monitored via alamar blue assay. GREB1 knockdown had no effect on the proliferation of T47D cells (Fig. 6B), suggesting constitutive Akt activation abrogates the need for GREB1 expression.

**Figure 6.**
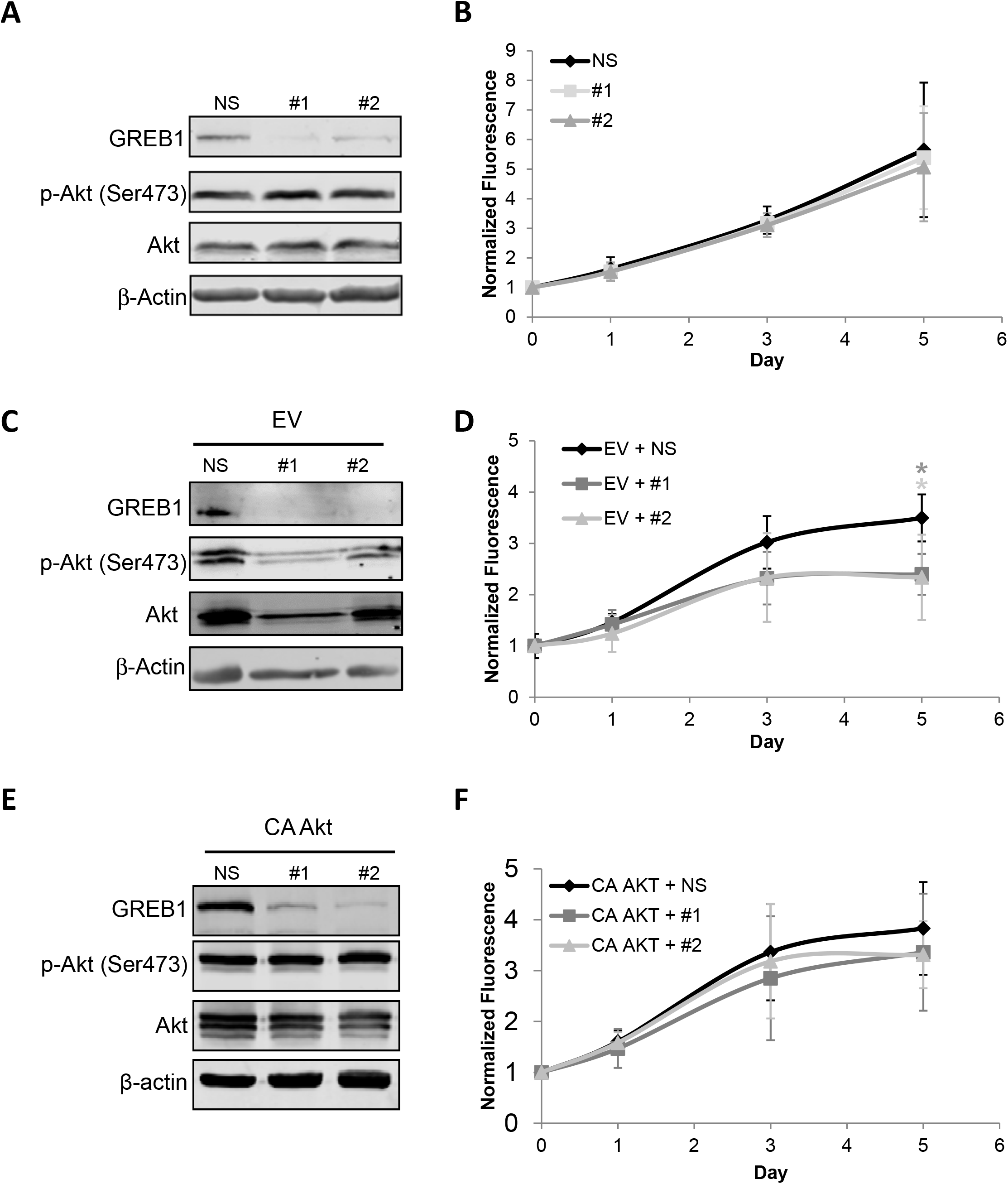
GREB1 regulates breast cancer proliferation through activation of the PI3K/Akt pathway. **A)** T47D cells were transduced with lentivirus expressing non-specific shRNA or shRNA targeted to GREB1 (shGREB1 #1 or shGREB1 #2). Immunoblot depicting the expression of indicated proteins. **B)** Proliferation was measured via alamar blue assay. Data are plotted as mean fluorescence normalized to Day 0 ± SD; n=3. **C)** MCF7 cells were transduced with lentivirus expressing empty vector (EV) and either non-specific shRNA or shRNA targeted to GREB1 (shGREB1 #1 or shGREB1 #2). Immunoblot showing the expression of labeled proteins. **D)** Proliferation was measured via almar blue assay. Data are plotted as mean fluorescence normalized to Day 0 ± SD; n=3. **E)** MCF7 cells were transduced with lentivirus expressing myristoylated Akt (CA AKT) and either non-specific shRNA or shRNA targeted to GREB1 (shGREB1 #1 or shGREB1 #2). Immunoblot demonstrating the expression of indicated protein. **F)** Proliferation was measured via alamar blue assay. Data are plotted as mean fluorescence normalized to Day 0 ± SD; n=3.

The proliferation of ER+ and GREB1-expressing MCF7 cells has previously been shown to be dependent on expression of GREB1 (16–18). While MCF7 cells also harbor a mutation in *PIK3CA*^E545K^, which is thought to be an activating mutation (34), these cells are still responsive to typical PI3K/Akt/mTOR-activating stimuli (Supplemental Fig. 1). Thus, we sought to determine if constitutively activated Akt (myristoylated-Akt) would rescue proliferation in GREB1-depleted MCF7 cells. MCF7 cells were transduced with empty vector lentivirus (EV) or lentivirus expressing constitutively activated Akt (CA AKT) in combination with lentivirus expressing shRNA targeted to a non-specific control (shNS) or to GREB1 (shGREB1 #1 or shGREB1 #2). Knockdown of endogenous GREB1 resulted in decreased Akt activation in cells co-transduced with GREB1-targeted shRNA and empty vector lentivirus (Fig. 6C) as well as parental cells infected with control lentivirus (Supplemental Fig. 4). The knockdown of GREB1 significantly impaired the growth of MCF7 cells co-transduced with empty vector lentivirus (Fig. 6D). However, expression of constitutively active Akt (Fig. 6E), rescues the proliferation phenotype caused by GREB1 knockdown to that of control transduced cells (Fig. 6F). Together, these data demonstrate that the primary mechanism by which GREB1 drives estrogen-dependent proliferation is through modulation of Akt activity. Interestingly, stable expression of constitutively active Akt results in long-term silencing of GREB1 (Supplemental Fig. 5), suggesting a feedback loop and potentially explaining the observation that GREB1 expression is reduced in hormone-refractory disease {Mohammed, 2013 #169}.

## Discussion

Despite extensive research on hormone signaling in breast cancer, the explicit mechanism by which ER drives proliferation remains largely undefined. In order to delineate this mechanism, concerted efforts have been made to identify ER-target genes involved in estrogen-induced proliferation of breast cancer cells. Several of these studies have identified GREB1 as a gene that is required for estrogen-stimulated proliferation of breast cancer cell lines (14, 16, 17). Previous studies have suggested that GREB1 regulates proliferation through modulation of ER activity (16). However, our findings show that GREB1 is not a potent regulator of ER activity and has the ability to affect the proliferation of breast cancer cell lines independent of ER expression and action (18). Here, we suggest a novel mechanism by which GREB1 regulates proliferation through fine-tuning of PI3K/Akt/mTOR signaling.

### GREB1 regulates proliferation of ER+ breast cancer cells through modulation of Akt activity

Several studies have indicated complex crosstalk between ER signaling and PI3K/Akt/mTOR pathway activation and implications for this crosstalk in resistance to endocrine therapy in breast cancer patients (7–9, 35–37). However, no studies have described a comprehensive connection between activation of these signaling pathways and proliferation of breast cancer cells. The difficulty to assess this connection is compounded by the fact that the vast majority of available breast cancer cell lines contain mutations involved in the PI3K/Akt/mTOR pathway (38). Although most ER+ breast cancer cell lines contain mutations within this pathway, the specific mutations have distinctly different effects on the activation of the PI3K/Akt/mTOR pathway. Specifically, breast cancer cell lines harboring *PIK3CA*^H1047R^ mutations (ex. T47D) have significantly higher intrinsic PI3K activity compared to breast cancer cell lines harboring *PIK3CA*^E545K^mutations (ex. MCF7), which have subtle effects on activation of the PI3K/Akt/mTOR pathway (11, 39). Using this to our advantage we show that in T47D cells, which harbor a constitutively active PI3K/Akt/mTOR pathway, GREB1 is no longer required for proliferation (Fig. 6A-B). However, in MCF7 cells, which harbor a *PIK3CA* mutation but still respond to pathway-activating stimuli, GREB1 is still required but knockdown of GREB1 can be rescued by constitutively active Akt (Fig. 6C-F). These data demonstrate that the requirement of GREB1 for hormone-responsive proliferation is dependent upon this novel function to alter Akt activity. Furthermore, the activation of the PI3K/Akt by GREB1 occurs through intracellular mechainisms (Fig. 4), highlighting a potentially new mode of modulating this signaling pathway without the need for RTK activation.

### GREB1 and endocrine resistance

Despite the clear association between GREB1 and proliferation of breast cancer cells, expression of GREB1 has been correlated to better prognosis in ER+ breast cancer patients (16, 40). In a study that included only patients that received adjuvant tamoxifen monotherapy, higher GREB1 expression correlated with both prolonged disease-free survival and sensitivity to tamoxifen treatment (40). Similarly, in an *in vitro* model of tamoxifen resistance, MCF7 cells that were resistant to tamoxifen treatment had significantly less GREB1 expression compared to the parental line, suggesting GREB1 expression is lost in hormone-refractory breast cancer cells (16). These data have implicated loss of GREB1 as a causal event for therapy-resistance. Here, we show that proliferation of breast cancer cells with constitutively active PI3K/Akt/mTOR signaling no longer require GREB1 expression (Fig. 6). Constitutive activation of the PI3K/Akt/mTOR pathway is frequently associated with resistance to endocrine therapies and is the basis for numerous clinical trials investigating PI3K/Akt/mTOR pathway inhibitors in endocrine-resistant patient populations (7–10). In patients with hormone-refractory disease with hyperactivation of the PI3K/Akt/mTOR pathway, the pressure to express GREB1 is lost. Thus, decreased GREB1 expression in advanced disease may be the result of constitutive PI3K/Akt/mTOR activity rather than a cause of therapeutic bypass. In support of this notion, stable expression of constitutively active Akt resulted in silencing of GREB1 expression after several passages (Supplemental Fig. 5). These findings warrant further research into the use of GREB1 as a clinical biomarker for treatment selection.

## Acknowledgements

We thank Paul Herman, Clarissa Wormsbaecher, Weiwei Liu, and Makanko Komara for critical reading of this manuscript prior to submission.

---

**Supplemental Figure S1.**
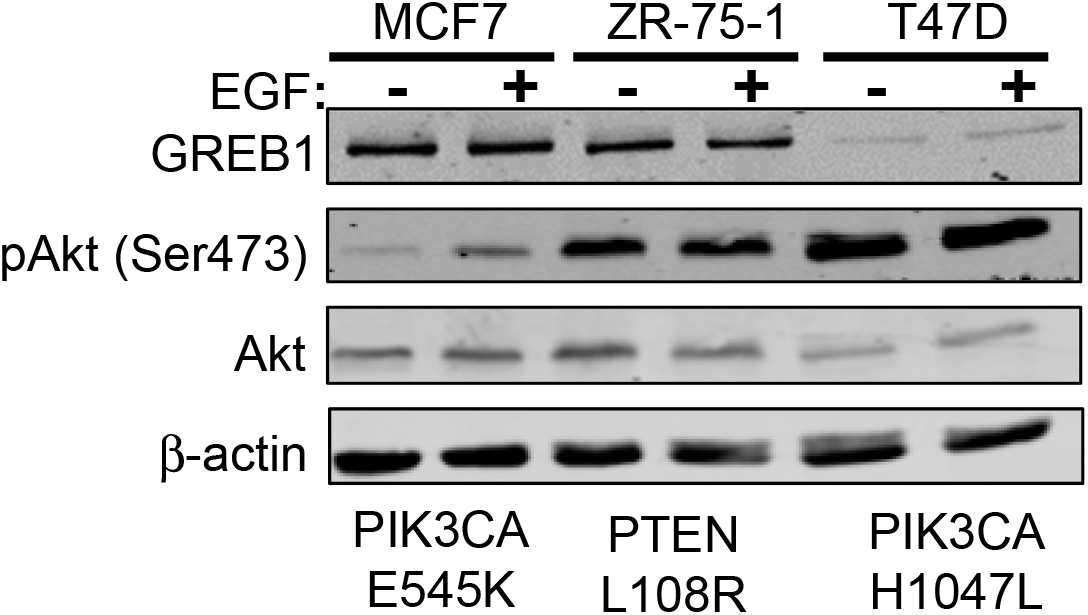
Different mutations in the PI3K pathway have varying levels of Akt activity in breast cancer cell lines. MCF7, ZR-75-1, and T47D cells were serum starved for 16 hours before stimulation with 1 ng/mL of EGF for 1 hour. Cell lysates were harvested and analyzed by immunoblot for the indicated proteins.

**Supplemental Figure S2.**
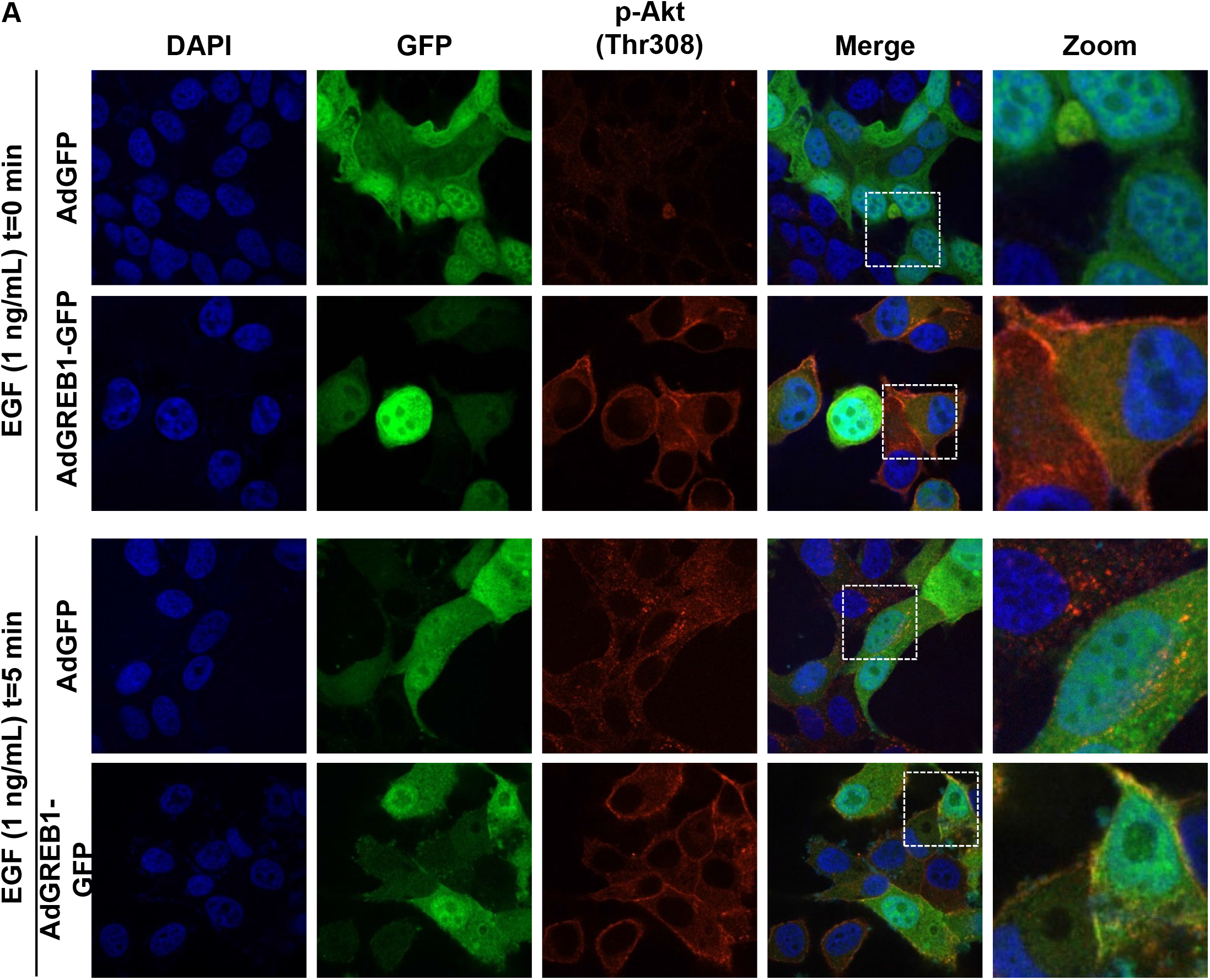
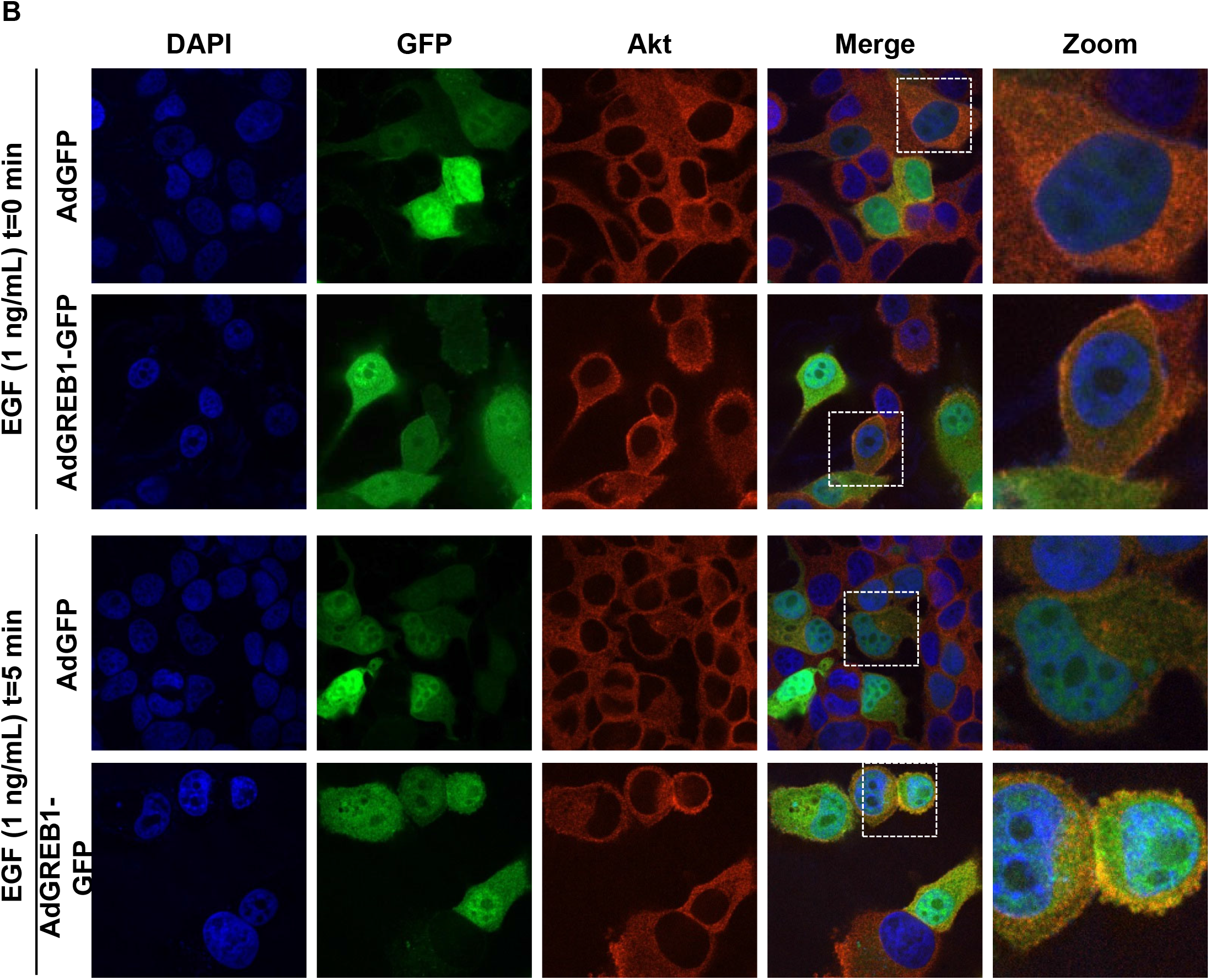
GREB1 overexpression induces Akt hyperactivation at the plasma membrane. MCF7 cells were transduced with adenovirus expressing GFP or GREB1. The cells were then cultured in serum-free media for 16 hours before being stimulated with 1 ng/mL EGF for 0 or 5 minutes. Cells were fixed and stained for DAPI and **A)** Akt or **B)** p-Akt (Thr308). Immunofluorescence microscopy was used to visualize the activation and localization of Akt.

**Supplemental Figure S3.**
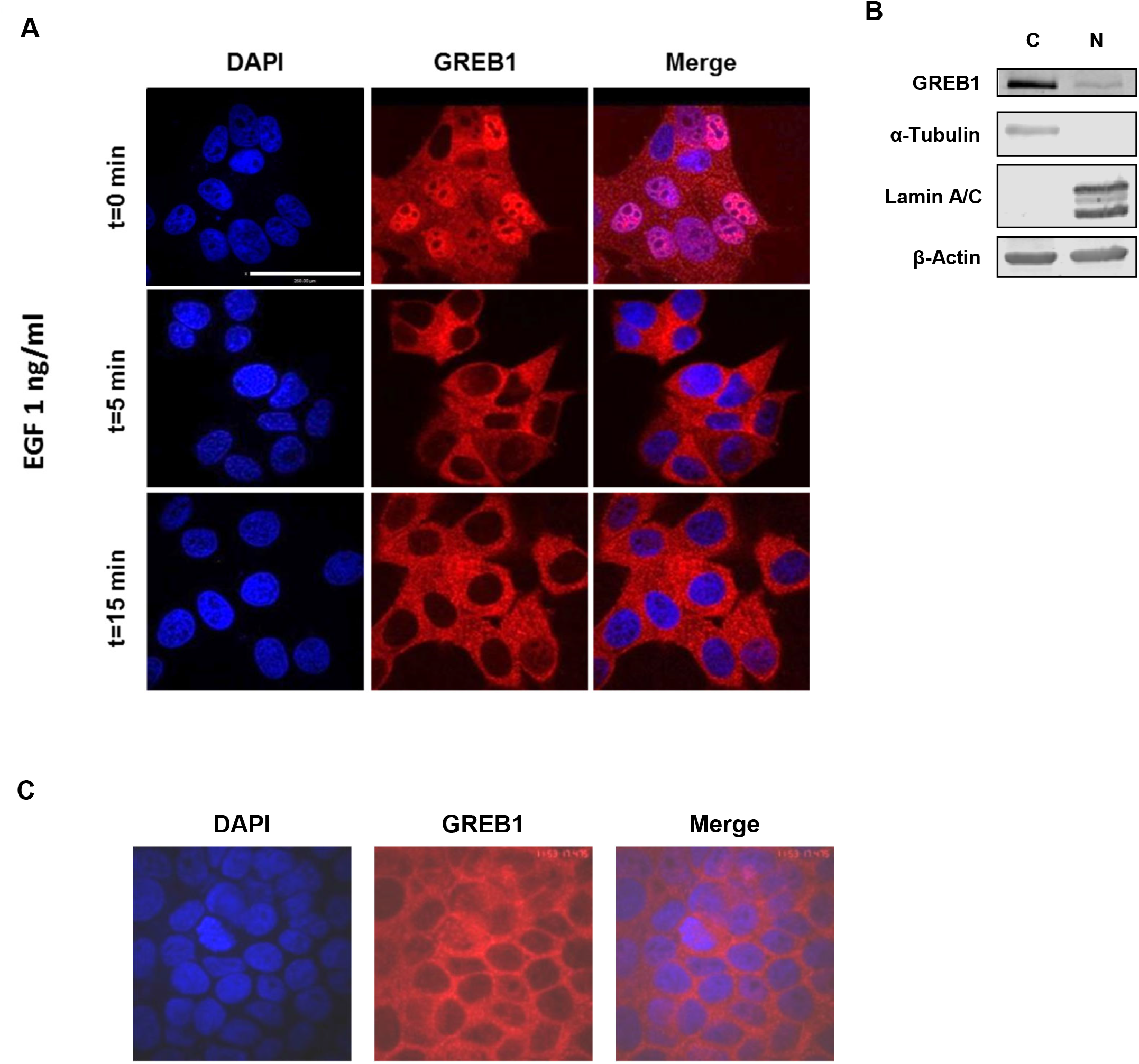
Endogenous GREB1 re-localizes to the cytoplasm under growth-stimulatory conditions. **A)** MCF7 cells were serum starved for 4 hours and stimulated with 1 ng/mL EGF for 0, 5, or 15 minutes. Cells were fixed and stained for DAPI and endogenous GREB1. Immunofluorescence microscopy was used to visualize GREB1 localization. **B)** Cytoplasmic and nuclear fractions were extracted from MCF7 whole cell lysate using high-speed centrifugation. Fractionated cell lysates were subjected to SDS-PAGE and analyzed via immunoblot for indicated proteins. **C)** MCF7 cells cultured in full serum media were fixed and stained for DAPI and endogenous GREB1. Immunofluorescence microscopy was used to visualize GREB1 localization under normal growth conditions.

**Supplemental Figure S4.**
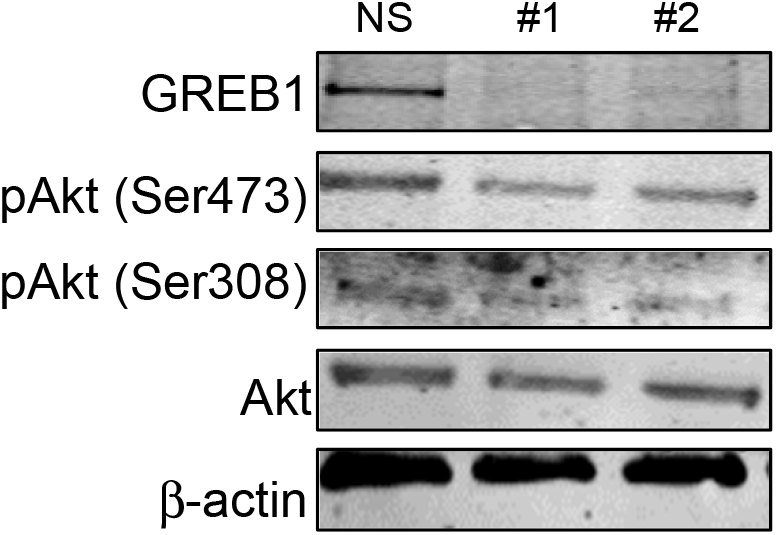
Endogenous GREB1 regulates Akt activation. MCF7 cells transduced with lentivirus expressing non-specific shRNA (shNS) or one of two shRNAs targeted to GREB1 (shGREB1 #1 or shGREB1 #2) were placed in serum/phenol red free media for 16 hours followed by 1 hour of activation with 1ng/ml EGF. Cells were harvested and lysates analyzed via immunoblot for Akt activation pathway.

**Supplemental Figure S5.**
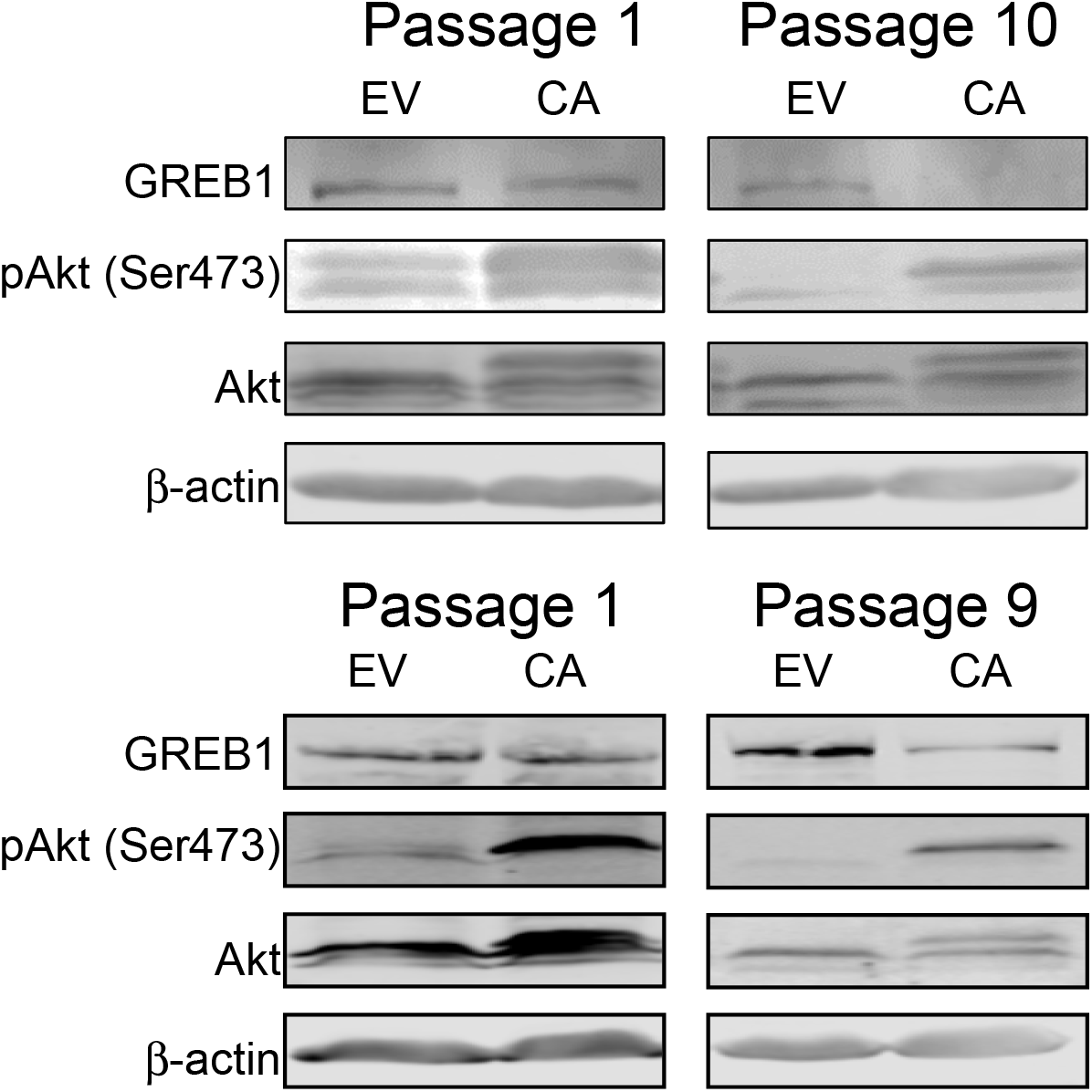
Constitutive Akt activation causes long-term silencing of GREB1 expression. Following transduction with control (EV) or constitutively active Akt (CA) expressing lentivirus, MCF7 stable lines were generated by placing cells on selection. Lysates from different passages were probed for GREB1 expression and Akt. activation Two distinct stable lines are depicted (top panel and bottom panel, respectively).

## References

1. Siegel RL, Miller KD, Jemal A. Cancer statistics, 2018. CA: a cancer journal for clinicians. 2018 Jan;68(1):7–30. PubMed PMID: 29313949. Epub 2018/01/10. eng.

2. Clarke R, Tyson JJ, Dixon JM. Endocrine resistance in breast cancer--An overview and update. Mol Cell Endocrinol. 2015 Dec 15;418 Pt 3:220–34. PubMed PMID: 26455641. Pubmed Central PMCID: PMC4684757.

3. Dixon JM. Endocrine Resistance in Breast Cancer. New Journal of Science. 2014;2014:1–27.

4. Libson S, Lippman M. A review of clinical aspects of breast cancer. International review of psychiatry (Abingdon, England). 2014 Feb;26(1):4–15. PubMed PMID: 24716497. Epub 2014/04/11. eng.

5. Patani N, Martin LA. Understanding response and resistance to oestrogen deprivation in ER-positive breast cancer. Mol Cell Endocrinol. 2014 Jan 25;382(1):683–94. PubMed PMID: 24121024. Epub 2013/10/15. eng.

6. Ali S, Coombes RC. Endocrine-responsive breast cancer and strategies for combating resistance. Nat Rev Cancer. 2002 Feb;2(2):101–12. PubMed PMID: 12635173.

7. Campbell RA, Bhat-Nakshatri P, Patel NM, Constantinidou D, Ali S, Nakshatri H. Phosphatidylinositol 3-kinase/AKT-mediated activation of estrogen receptor alpha: a new model for anti-estrogen resistance. J Biol Chem. 2001 Mar 30;276(13):9817–24. PubMed PMID: 11139588. Epub 2001/01/15. eng.

8. Miller TW, Balko JM, Arteaga CL. Phosphatidylinositol 3-kinase and antiestrogen resistance in breast cancer. Journal of clinical oncology: official journal of the American Society of Clinical Oncology. 2011 Nov 20;29(33):4452–61. PubMed PMID: 22010023. Pubmed Central PMCID: PMC3221526. Epub 2011/10/20. eng.

9. Miller TW, Hennessy BT, Gonzalez-Angulo AM, Fox EM, Mills GB, Chen H, et al. Hyperactivation of phosphatidylinositol-3 kinase promotes escape from hormone dependence in estrogen receptor-positive human breast cancer. The Journal of clinical investigation. 2010 Jul;120(7):2406–13. PubMed PMID: 20530877. Pubmed Central PMCID: PMC2898598. Epub 2010/06/10. eng.

10. Miller WR, Larionov A, Renshaw L, Anderson TJ, Walker JR, Krause A, et al. Gene expression profiles differentiating between breast cancers clinically responsive or resistant to letrozole. Journal of clinical oncology: official journal of the American Society of Clinical Oncology. 2009 Mar 20;27(9):1382–7. PubMed PMID: 19224856. Epub 2009/02/20. eng.

11. Ellis MJ, Perou CM. The genomic landscape of breast cancer as a therapeutic roadmap. Cancer discovery. 2013 Jan;3(1):27–34. PubMed PMID: 23319768. Pubmed Central PMCID: PMC3553590. Epub 2013/01/16. eng.

12. Simoncini T, Hafezi-Moghadam A, Brazil DP, Ley K, Chin WW, Liao JK. Interaction of oestrogen receptor with the regulatory subunit of phosphatidylinositol-3-OH kinase. Nature. 2000 Sep 28;407(6803):538–41. PubMed PMID: 11029009. Pubmed Central PMCID: PMC2670482. Epub 2000/10/12. eng.

13. Creighton CJ, Fu X, Hennessy BT, Casa AJ, Zhang Y, Gonzalez-Angulo AM, et al. Proteomic and transcriptomic profiling reveals a link between the PI3K pathway and lower estrogen-receptor (ER) levels and activity in ER+ breast cancer. Breast cancer research: BCR. 2010;12(3):R40. PubMed PMID: 20569503. Pubmed Central PMCID: PMC2917035. Epub 2010/06/24. eng.

14. Ghosh MG, Thompson DA, Weigel RJ. PDZK1 and GREB1 are estrogen-regulated genes expressed in hormone-responsive breast cancer. Cancer Res. 2000 Nov 15;60(22):6367–75. PubMed PMID: 11103799. Epub 2000/12/05. eng.

15. Hnatyszyn HJ, Liu M, Hilger A, Herbert L, Gomez-Fernandez CR, Jorda M, et al. Correlation of GREB1 mRNA with protein expression in breast cancer: validation of a novel GREB1 monoclonal antibody. Breast Cancer Res Treat. 2010 Jul;122(2):371–80. PubMed PMID: 19842031.

16. Mohammed H, D’Santos C, Serandour AA, Ali HR, Brown GD, Atkins A, et al. Endogenous purification reveals GREB1 as a key estrogen receptor regulatory factor. Cell Rep. 2013 Feb 21;3(2):342–9. PubMed PMID: 23403292.

17. Rae JM, Johnson MD, Scheys JO, Cordero KE, Larios JM, Lippman ME. GREB 1 is a critical regulator of hormone dependent breast cancer growth. Breast Cancer Res Treat. 2005 Jul;92(2):141–9. PubMed PMID: 15986123.

18. Haines CN, Braunreiter KM, Mo XM, Burd CJ. GREB1 isoforms regulate proliferation independent of ERalpha co-regulator activities in breast cancer. Endocrine-related cancer. 2018 Jul;25(7):735–46. PubMed PMID: 29695586. Epub 2018/04/27. eng.

19. Burd CJ, Petre CE, Moghadam H, Wilson EM, Knudsen KE. Cyclin D1 binding to the androgen receptor (AR) NH2-terminal domain inhibits activation function 2 association and reveals dual roles for AR corepression. Mol Endocrinol. 2005 Mar;19(3):607–20. PubMed PMID: 15539430. Epub 2004/11/13. eng.

20. Lee MS, Jeong MH, Lee HW, Han HJ, Ko A, Hewitt SM, et al. PI3K/AKT activation induces PTEN ubiquitination and destabilization accelerating tumourigenesis. Nature communications. 2015 Jul 17;6:7769. PubMed PMID: 26183061. Pubmed Central PMCID: PMC4518267. Epub 2015/07/18. eng.

21. Boehm JS, Zhao JJ, Yao J, Kim SY, Firestein R, Dunn IF, et al. Integrative genomic approaches identify IKBKE as a breast cancer oncogene. Cell. 2007 Jun 15;129(6):1065–79. PubMed PMID: 17574021. Epub 2007/06/19. eng.

22. Patterson AR, Mo X, Shapiro A, Wernke KE, Archer TK, Burd CJ. Sustained reprogramming of the estrogen response after chronic exposure to endocrine disruptors. Mol Endocrinol. 2015 Mar;29(3):384–95. PubMed PMID: 25594248. Pubmed Central PMCID: PMC4347288. Epub 2015/01/17. eng.

23. Guan X, LaPak KM, Hennessey RC, Yu CY, Shakya R, Zhang J, et al. Stromal Senescence By Prolonged CDK4/6 Inhibition Potentiates Tumor Growth. Molecular cancer research: MCR. 2017 Mar;15(3):237–49. PubMed PMID: 28039358. Pubmed Central PMCID: 5334447.

24. Debacq-Chainiaux F, Erusalimsky JD, Campisi J, Toussaint O. Protocols to detect senescence-associated beta-galactosidase (SA-betagal) activity, a biomarker of senescent cells in culture and in vivo. Nat Protoc. 2009;4(12):1798–806. PubMed PMID: 20010931. Epub 2009/12/17. eng.

25. Xu Y, Li N, Xiang R, Sun P. Emerging roles of the p38 MAPK and PI3K/AKT/mTOR pathways in oncogene-induced senescence. Trends Biochem Sci. 2014 Jun;39(6):268–76. PubMed PMID: 24818748. Pubmed Central PMCID: PMC4358807.

26. Freund A, Patil CK, Campisi J. p38MAPK is a novel DNA damage response-independent regulator of the senescence-associated secretory phenotype. EMBO J. 2011 Apr 20;30(8):1536–48. PubMed PMID: 21399611. Pubmed Central PMCID: PMC3102277.

27. Courtois-Cox S, Jones SL, Cichowski K. Many roads lead to oncogene-induced senescence. Oncogene. 2008 May 01;27(20):2801–9. PubMed PMID: 18193093.

28. She QB, Chandarlapaty S, Ye Q, Lobo J, Haskell KM, Leander KR, et al. Breast tumor cells with PI3K mutation or HER2 amplification are selectively addicted to Akt signaling. PLoS One. 2008 Aug 26;3(8):e3065. PubMed PMID: 18725974. Pubmed Central PMCID: PMC2516933. Epub 2008/08/30. eng.

29. Manning BD, Toker A. AKT/PKB Signaling: Navigating the Network. Cell. 2017 Apr 20;169(3):381–405. PubMed PMID: 28431241. Pubmed Central PMCID: PMC5546324. Epub 2017/04/22. eng.

30. Du K, Tsichlis PN. Regulation of the Akt kinase by interacting proteins. Oncogene. 2005 Nov 14;24(50):7401–9. PubMed PMID: 16288287. Epub 2005/11/17. eng.

31. Bellacosa A, Chan TO, Ahmed NN, Datta K, Malstrom S, Stokoe D, et al. Akt activation by growth factors is a multiple-step process: the role of the PH domain. Oncogene. 1998 Jul 23;17(3):313–25. PubMed PMID: 9690513. Epub 1998/08/05. eng.

32. Bellacosa A, Kumar CC, Di Cristofano A, Testa JR. Activation of AKT kinases in cancer: implications for therapeutic targeting. Advances in cancer research. 2005;94:29–86. PubMed PMID: 16095999. Epub 2005/08/13. eng.

33. Liu P, Cheng H, Roberts TM, Zhao JJ. Targeting the phosphoinositide 3-kinase pathway in cancer. Nature reviews Drug discovery. 2009;8(8):627–44. PubMed PMID: 19644473.

34. Bosch A, Li Z, Bergamaschi A, Ellis H, Toska E, Prat A, et al. PI3K inhibition results in enhanced estrogen receptor function and dependence in hormone receptor-positive breast cancer. Science translational medicine. 2015 Apr 15;7(283):283ra51. PubMed PMID: 25877889. Pubmed Central PMCID: PMC4433148. Epub 2015/04/17. eng.

35. Bhat-Nakshatri P, Wang G, Appaiah H, Luktuke N, Carroll JS, Geistlinger TR, et al. AKT alters genome-wide estrogen receptor alpha binding and impacts estrogen signaling in breast cancer. Mol Cell Biol. 2008 Dec;28(24):7487–503. PubMed PMID: 18838536. Pubmed Central PMCID: PMC2593438. Epub 2008/10/08. eng.

36. Chen D, Washbrook E, Sarwar N, Bates GJ, Pace PE, Thirunuvakkarasu V, et al. Phosphorylation of human estrogen receptor alpha at serine 118 by two distinct signal transduction pathways revealed by phosphorylation-specific antisera. Oncogene. 2002 Jul 25;21(32):4921–31. PubMed PMID: 12118371. Epub 2002/07/16. eng.

37. Kato S, Endoh H, Masuhiro Y, Kitamoto T, Uchiyama S, Sasaki H, et al. Activation of the estrogen receptor through phosphorylation by mitogen-activated protein kinase. Science (New York, NY). 1995 Dec 1;270(5241):1491–4. PubMed PMID: 7491495. Epub 1995/12/01. eng.

38. Smith SE, Mellor P, Ward AK, Kendall S, McDonald M, Vizeacoumar FS, et al. Molecular characterization of breast cancer cell lines through multiple omic approaches. Breast cancer research: BCR. 2017 Jun 5;19(1):65. PubMed PMID: 28583138. Pubmed Central PMCID: PMC5460504. Epub 2017/06/07. eng.

39. Jamieson S, Flanagan JU, Kolekar S, Buchanan C, Kendall JD, Lee WJ, et al. A drug targeting only p110alpha can block phosphoinositide 3-kinase signalling and tumour growth in certain cell types. Biochem J. 2011 Aug 15;438(1):53–62. PubMed PMID: 21668414. Pubmed Central PMCID: PMC3174055. Epub 2011/06/15. eng.

40. Wu Y, Zhang Z, Cenciarini ME, Proietti CJ, Amasino M, Hong T, et al. Tamoxifen Resistance in Breast Cancer Is Regulated by the EZH2-ERalpha-GREB1 Transcriptional Axis. Cancer Res. 2018 Feb 1;78(3):671–84. PubMed PMID: 29212856. Pubmed Central PMCID: PMC5967248. Epub 2017/12/08. eng.

